# Loss of Bardet-Biedl syndrome proteins causes synaptic aberrations in principal neurons

**DOI:** 10.1101/580399

**Authors:** Naila Haq, Christoph Schmidt-Hieber, Fernando J. Sialana, Lorenza Ciani, Janosch P. Heller, Michelle Stewart, Liz Bentley, Sara Wells, Richard J. Rodenburg, Patrick M Nolan, Elizabeth Forsythe, Michael C. Wu, Gert Lubec, Michael Häusser, Philip L. Beales, Sofia Christou-Savina

**Affiliations:** Great Ormond Street Institute of Child Health, University College London, UK; Institut Pasteur, Paris, France; Department of Pharmaceutical Chemistry, University of Vienna, Austria; Department of Cell and Developmental Biology, University College London, UK; Institute of Neurology, University College London, UK; Mary Lyon Centre, MRC Harwell, UK; Radboud Center for Mitochondrial Medicine, Translational Metabolic Laboratory, Department of Pediatrics, Radboud University Medical Centre, Nijmegen, The Netherlands; Neurodigitech, LLC, San Diego, US; Wolfson Institute for Biomedical Research, University College London, UK

## Abstract

Bardet-Biedl syndrome (BBS), a ciliopathy, is a rare genetic condition characterised by retina degeneration, obesity, kidney failure and cognitive impairment. In spite of a progress made in general understanding of BBS aetiology, the molecular mechanism of cognitive impairment remains elusive. Here we report that loss of Bardet-Biedl syndrome proteins causes synaptic dysfunction in principal neurons providing possible explanation for cognitive impairment phenotype in BBS patients. Using synaptosomal proteomics and immunocytochemistry we demonstrate the presence of Bbs in postsynaptic density of hippocampal neurons. Loss of Bbs results in the significant reduction of dendritic spines in principal neurons of *Bbs* mice models. Furthermore, we demonstrate that spine deficiency correlates with events that destabilize spine architecture, such as, impaired spine membrane receptors signalling known to be involved in the maintenance of dendritic spines. Finally, we show that voluntary exercise rescues spine deficiency in the neurons. Based on our data, we propose a model in which Bbs proteins, similar to their function in primary cilia, regulate trafficking of signalling receptors into and out of the membrane of dendritic spines, thus providing the basis for synaptic plasticity.

## Introduction

Dendritic spines are small protrusions that cover the dendrites of most principal neurons in the vertebrate central nervous system (CNS), where they typically serve as the postsynaptic part of excitatory synapses (1). Recent studies have revealed that alterations in dendritic spines are associated with a wide range of cognitive related conditions ranging from rare monogenic neurodevelopmental syndromes to common psychiatric diseases, including schizophrenia and bipolar disorder (2–4). Dendritic spine shape, size and number are regulated in a spatiotemporal manner that is tightly coordinated with synaptic function and plasticity (5, 6). Formation, maintenance, stability and pruning of spines is tightly coordinated by a wide range of surface receptors that, when activated by extracellular ligands, trigger diverse downstream signalling pathways. Membrane receptors such as Insulin-Like Growth Factor (IGF-1R), ephrin B (EphB), and TrkB have profound effects on neuroplasticity in CNS (7, 8). There are myriad of ways in which the activation of these receptors may mediate neuroplasticity including modulation of Akt/mTOR pathway, macroautophagy, small GTPases activity, and glutamate receptors (GluRs) membrane expression (9).

BBS proteins comprise of a group of unrelated ciliary proteins that, when mutated, cause a rare genetic disorder, Bardet-Biedl syndrome (BBS). BBS is a genetically heterogeneous, autosomal recessive disorder characterised by early-onset retinal degeneration, obesity, polydactyly, renal dysfunction, and cognitive impairment (10, 11). BBS was one of the first multi-system disorder ascribed to dysfunctional non-motile cilia, microtubule based membranous projections protruding from the cell surface of most mammalian cells, including neurons (12). It was shown that eight highly conserved BBS proteins form a coat like complex, BBSome that is responsible for the trafficking of signalling receptors into and out of cilia (13). Loss of BBS proteins affect entry and exit of signalling receptors such as somatostatin receptor type 3 (Sstr3), melanin-concentrating hormone receptor 1 (Mchr1), dopamine receptors (D1), Gpr161, and BDNF receptors (TrkB) (14, 15). One area of growing interest is the role of BBS proteins in cognition. The majority of individuals with BBS, experience developmental disabilities ranging from mild impairment or delayed emotional development to severe mental and psychiatric disorders (16). The frequency of neuro-psychiatric disorders and autism in BBS group exceeds the incidence rate of these disorders in the general population (17).

Here we show for the first time significant morphological aberrations of dendritic spines in the principal neurons of *Bbs* murine models which correlate with impaired contextual and cued fear conditioning test. While investigating the molecular mechanisms of dendritic spine loss in *Bbs* mice models, we excluded reduced synaptic activity and mitochondria dysfunction as a likely cause of spine loss. At the same time we found that spine deficiency correlates with impaired spine membrane receptors signalling known to be involved in the maintenance of dendritic spines among them IGF-1R and ephrin B receptor signalling. Moreover, our finding of BBS proteins localization to the postsynaptic densities (PSD) of hippocampal neurons prompted us to propose a model for BBS proteins function in synapses. Together, our data reveal that BBS proteins, their role so far being confined only to the functional maintenance of cilia, play an important role in synaptic structure and function and suggest that aberrant spine formation and maintenance may underlie the cognitive impairment in BBS patients.

## Results

### Loss of Bbs proteins affect principal neuron morphology

To investigate the effect of Bbs protein loss on the neuronal structure we analysed the dendritic morphology of principal neurons of Bbs mice models which closely mimic the human BBS phenotype. We measured dendritic length, spine count and spine density of dentate gyrus (DG), basolateral amygdala (BLA) and layer V pyramidal neurons of the frontal cortex using a Golgi-Cox impregnation method. We found that the total spine density was reduced by 55% in DG granule cells of 6 weeks old *Bbs4^−/−^* mice (Fig.1a,b and Supplementary Movie 1). Total basal and apical spine density of layer V neurons was reduced by 55% and 54%, (Fig.1a,e), respectively, and total basal and apical spine density of BLA neurons was reduced by 23% and 22%, respectively (Fig.1a,j). Sholl analysis revealed a significant reduction in spine density of all branches and per 30 μm interval in DG with the exception of the most distal branch and 300 μm circle in *Bbs4^−/−^* mice (Fig.1c,d). Similar Sholl analysis results were found in apical and basal dendrites of Layer V neurons where dendritic spine density per branch order and per 30 μm interval was affected (Fig.1f-i). Apical and basal BLA dendrites of *Bbs4* knockout mice revealed unequal pattern in spine reduction affecting only few branches and concentric circles (Fig.1k-n). A number of dendritic intersections in DG, BLA and Layer V neurons were not affected when compared to control mice (Supplementary Fig.1 a-e). Total dendritic length was reduced by 48% in DG neurons and by 25% and 12% in basal and apical dendrites of layer V cortex neurons. BLA apical and basal dendrites showed a statistically significant length reduction of 14% and 19%, respectively (Supplementary Fig.1f-j). Overall, this data show significant aberrations in dendritic morphology.

**Fig.1.**
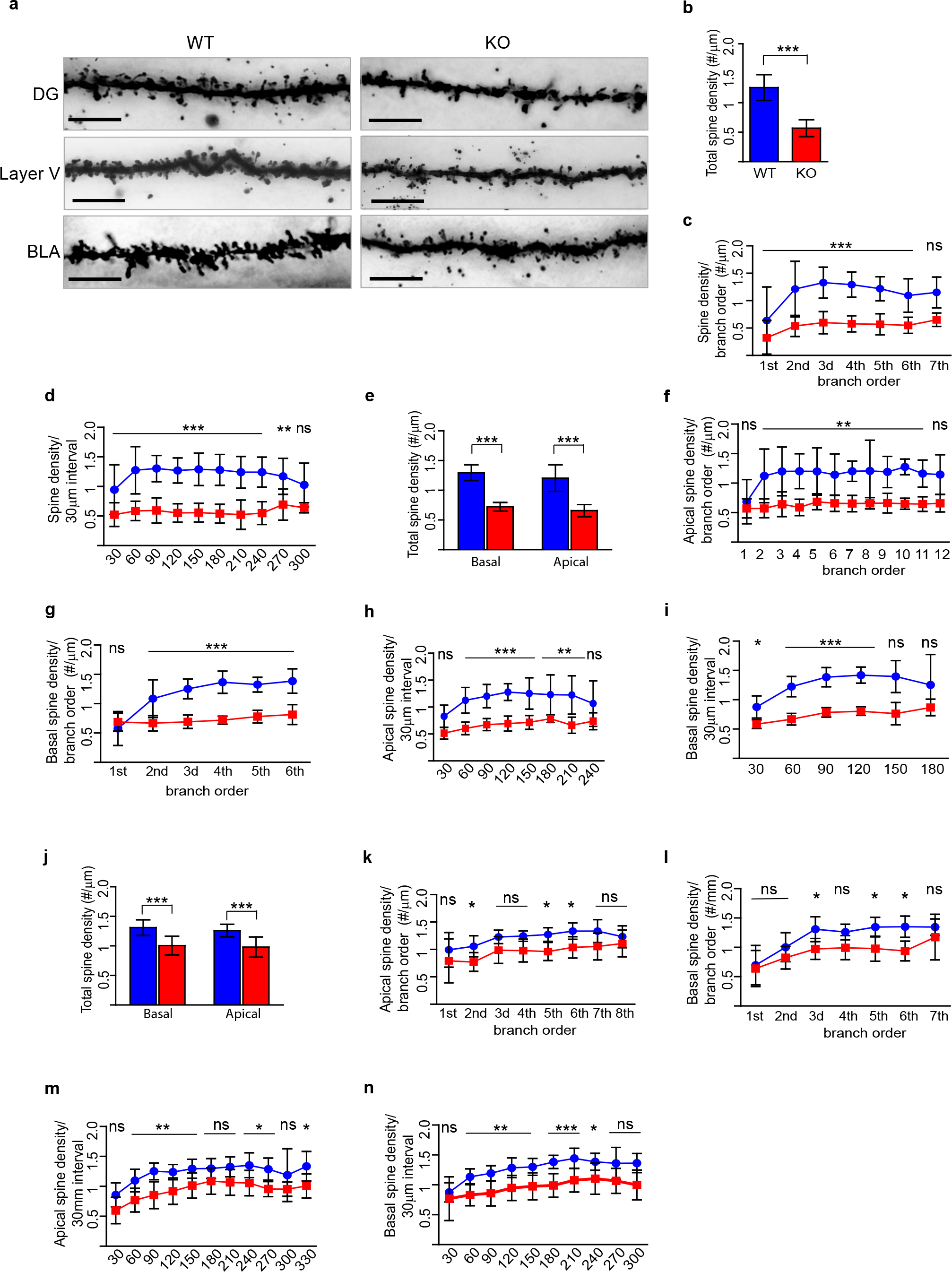
Dendritic Spine morphology of Dentate Gyrus (DG), Basolateral Amygdala (BLA) and Frontal Cortex neurons is affected by the loss of Bbs4 protein. **(a)** Representative images of Golgi-impregnated dentate granule (DG), Basolateral Amygdala (BLA) and Layer V pyramidal neurons of 6 weeks old *Bbs4*^−/−^ and *Bbs4*^+/+^ mice at the age of 6 weeks (100x; scale bar 5 μm) **(b-d)** Total spine density of DG neurons **(b)**, Spine density per branch order **(c)**, Spine density per 30 μm interval **(d)** **(e-i)** Total spine density of layer V pyramidal neurons **(e)**, Spine density in apical dendrites per branch order **(f)**, Spine density in basal dendrites per branch order **(g)**, Spine density in apical dendrites per 30 μm interval **(h)**, Spine density in basal dendrites per 30 μm interval **(i)** **(j-n)** Total spine density of basolateral amygdala neurons **(j)**, Spine density in apical dendrites per branch order **(k)**, Spine density in basal dendrites per branch order **(i)**, Spine density in apical dendrites per 30 μm interval **(m)**, Spine density in basal dendrites per 30 μm interval **(n)** (N_mice/WT_=5; N_mice/KO_=7, N_cells/WT_=25, N_cells /KO_=35, Mean ± SD, ***P<0.001; **P<0.01; *P<0.05). One-way *ANOVA*, Tukey *post hoc* test except for b, e, j where unpaired *t-*test was used

In order to investigate when the dendritic architecture of *Bbs4^−/−^* DG neurons begins to change, we analysed dendritic length, spine density per branch order and per 30μm intervals at P21 mice. The results were similar to those obtained at 6 weeks of age: a significant reduction in dendritic length, spine density per branch order, and per 30μm interval (Supplementary Fig.2a, g-k). However, at E19.5, the density of filopodia in *Bbs4^−/−^* DG neurons was not affected (Supplementary Fig.2a-f). Taken together, the *Bbs4^−/−^* murine model show a progressive decrease in dendritic spine density at 3 weeks old mice (38%) and 6 weeks old mice (55%) but not during late embryonic stages (Supplementary Fig.2l).

To investigate whether similar dendritic abnormalities can be detected in the other *Bbs* models, we analysed DG dendrite morphology of 3 weeks old *Bbs5* knockout and *Bbs1* M390R knock-in models. Notably, loss of Bbs5 protein led to significant reduction in DG dendritic length (34%) and overall spine density (32%) in the dentate granule cells of knockout mice (Supplementary Fig.3a-c). Sholl analysis also revealed abnormal spine density in *Bbs5*^−/−^mice with a significant spine reduction from the 2^nd^ to 5^th^ branch order and from 60μm to 150μm interval, respectively (Supplementary Fig.3d-f). Interestingly, the *Bbs1 M390R* knock-in model was associated with consistent but moderate abnormalities in spino- and dendritogenesis of DG neurons showing only a 10% reduction in the total spine density (Supplementary Fig.3a, g-k). This finding is in agreement with our clinical observations that *BBS1 M390R* patients have the mildest cognitive phenotype.

Dendritic spines, especially on pyramidal cells, could be subdivided into categories based on their size and shape (Supplementary Fig.4a) (18). We next asked whether certain subtypes of spines were over-represented on DG neurons of *Bbs4^−/−^* mice. We observed that total spine count was reduced in all spine subtypes except ‘branched’ spines, however, when the reduction of dendritic length of *Bbs4* neurons were taken into account, we found that only the density of ‘thin’ spines was significantly reduced (22%) (Supplementary Fig.4b-e).

### Reduced contextual and cued fear memory but no impairment in anxiety-like behaviour in *Bbs4*^−/−^ mice

Hippocampus, amygdala and prefrontal cortex are structures involved in learning, memory and social interaction. To investigate whether the loss of dendritic spines of DG, BLA and prefrontal cortex neurons correlates with behavioural changes of the *Bbs4^−/−^* mice, we performed analysis of data obtained from a battery of behavioural tests in 8 weeks old knockout and control mice. The contextual and cued fear conditioning tests are widely used to assess fear memory. In the conditioning session at Day 1, freezing behaviour and distance travelled during the first 150 s without introducing a conditioned stimulus (tone) and unconditioned stimulus (footshock) were used to evaluate baseline activity in the novel environment in the context fear experiment. There was no effect of Bbs4 loss on the percentage of freezing or distance travelled in baseline activity of *Bbs4^−/−^* males and female mice. However, after introduction of the paired tone-foot shock stimulus at Day 1, *post-hoc* analysis revealed a significant decrease in percentage of freezing and an increase in the distance travelled on Day 2 of *Bbs4^−/−^* female mice compared to female wild type mice. Male mice showed a significant decrease in percentage of freezing and an increase in distance travelled when the un-paired, two-tailed *t* test (p<0.05) was used (Fig.2a, b, d). Altered context at Day 2 before the introduction of the tone was used as a baseline for the cue data. There was no significant change in the percentage of freezing in altered context baseline activity (Fig.2c, d). During the cued conditioning session with the tone (Day 2, 180-360 s), the percentage of freezing was significantly reduced and distance travelled increased in *Bbs4^−/−^* males (Fig.2c, d). This set of results indicates that loss of Bbs4 protein affects contextual and cued fear memory.

**Fig.2.**
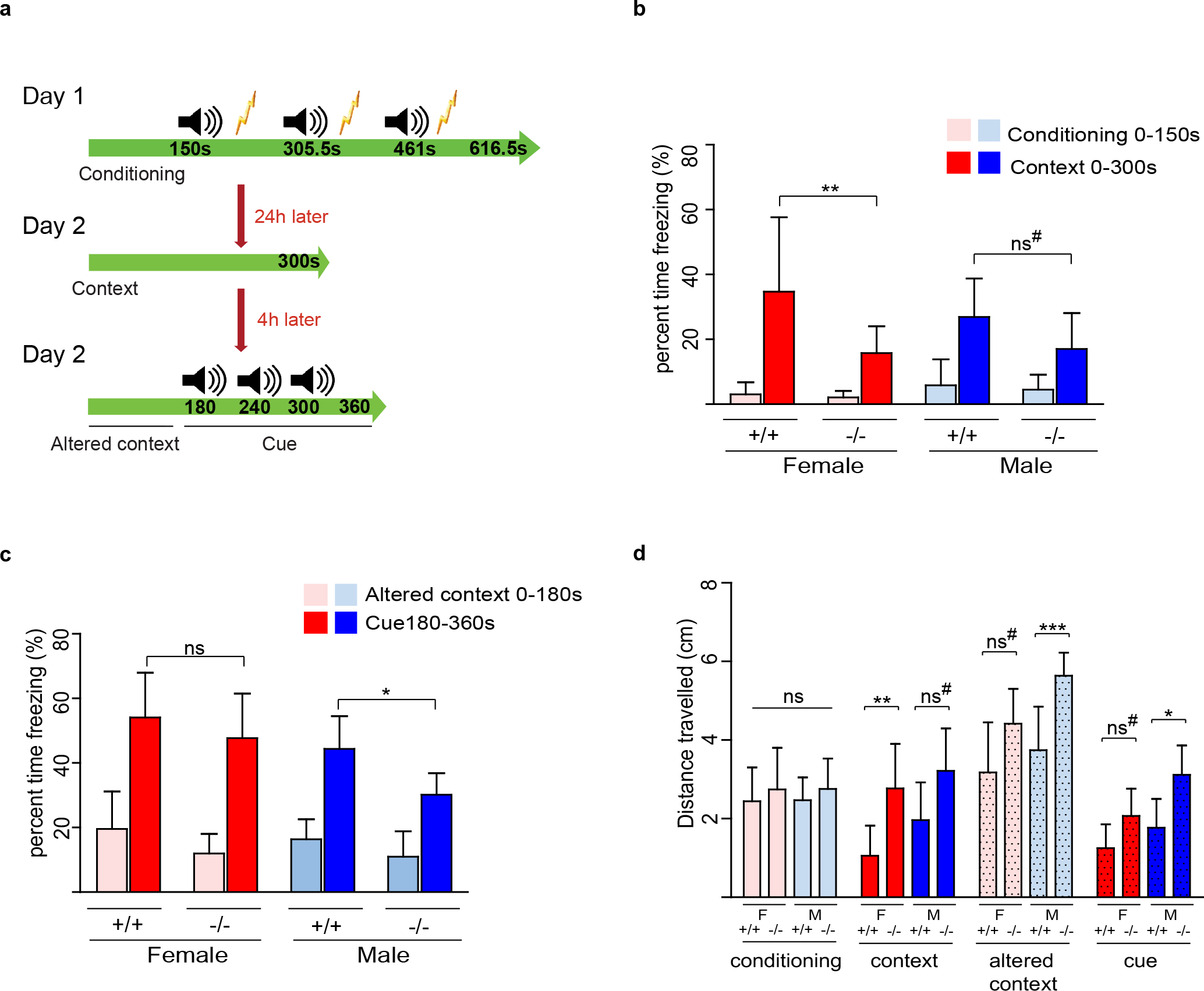
Aberrant fear conditioning behaviour in *Bbs4* knockout mice. **(a)** Schematic presentation of the contextual and cued fear conditioning test. At Day 1 mice were placed in fear conditioning chamber for 616.6 s. After 150 s, a 5 s tone is played, followed by a 0.5 s, 0.5 mA shock. The tone and shock are repeated two more times at 150 s interval. At Day 2 mice were paced in exactly the same chamber for 300 s without tones or shocks. After 4 hours (Day 2) mice are placed in altered context and left for 180 s. At 180 s a 5 s tone is played which is repeated twice at 60 s intervals. First 150 s of the conditioning trial were used as a baseline for the context data. First 180 s in the altered context were used as the base line for the cue data. **(b)** Freezing (%) in the contextual memory test **(c)** Freezing (%) in the cue memory test **(d)** Distance travelled (cm) in the conditioning, context test, and cued test (Females: N_WT_=11, N_KO_=11; Males: N_WT_=13, N_KO_=12; Mean ± SD, ***P<0.001; **P<0.01; *P<0.05). One-way *ANOVA*, Tukey post hoc test. ^#^ It was noted that there was a significant level of reduction of percent time freezing and distance travelled in *Bbs4* knockout mice when un-paired, two-tailed *t* test (p<0.05) was used

Anxiety-like behaviour, locomotor activity and spatial memory were also assessed by using open field, light dark box and paddling Y-maze. No significant changes were obtained between *Bbs4^+/+^* and *Bbs4^−/−^*mice (data not shown).

### Miniature excitatory postsynaptic currents (mEPSCs) amplitude is increased in *Bbs4*^−/−^ DG neurons

The morphology of dendritic spines is highly dynamic and their formation and maintenance depend on neuronal activity (19). As all existing *Bbs* mice models exhibit various age dependent sensory deficits such as loss of vision, obesity and thermo- and mechanosensory aberrations (11, 20) we measured intrinsic properties and synaptic function of hippocampal granule cells of 6 weeks old *Bbs4^−/−^* and *Bbs4^+/+^* mice *in vitro*. We found that intrinsic properties of *Bbs4^−/−^* neurons were unaffected (Fig.3a-c). To evaluate the synaptic properties of granule cells in these two groups, we measured mEPSCs. Notably, while the frequency of mEPSCs was not different between the two groups, mEPSC amplitudes were significantly larger in *Bbs4^−/−^* neurons (Fig.3d-f). These data show that there is no reduction in the brain activity in *Bbs4^−/−^* mice which makes it unlikely that reduced neuronal activity might be the cause of reduced spine density. It also suggests an activation of compensatory mechanisms at the presynaptic and/or postsynaptic sites in response to spine loss.

**Fig.3.**
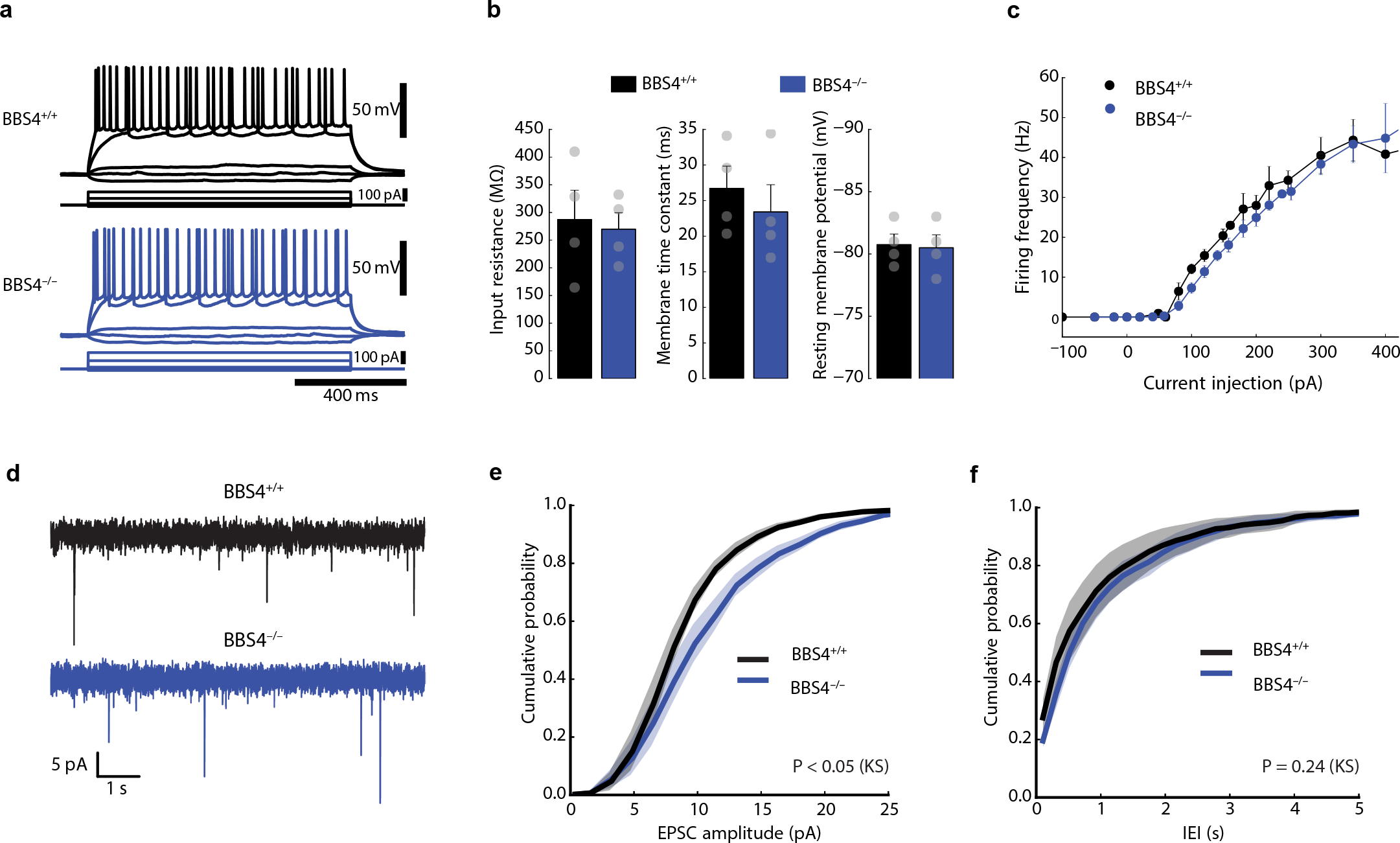
Comparison of electrophysiological properties of hippocampal neurons between *Bbs4* knockout and control mice *in vitro*. **(a)** Comparison of firing patterns in response to current injections during current clamp recordings from hippocampal granule cells **(b)** Bar graphs summarizing passive membrane properties. No significant differences were found between *Bbs4*^−/−^ and *Bbs4*^+/+^ mice in input resistance (left), membrane time constant (middle) and resting membrane potential (right) **(c)** Firing frequency was plotted against current injection amplitudes. No significant differences were found between *Bbs4*^−/−^ and *Bbs4*^+/+^ mice **(d)** Miniature EPSCs were recorded from hippocampal granule cells in *Bbs4*^−/−^ and *Bbs4*^+/+^ mice (N=6) **(e)** Cumulative probability plot comparing mEPSC amplitudes between *Bbs4*^−/−^ and *Bbs4*+/+ mice. mEPSC amplitudes in granule cells of *Bbs4*^−/−^ mice are significantly larger (N= 6, P < 0.05, Kolmogorov-Smirnov test) **(f)** Cumulative probability plot comparing inter-event intervals (IEI) of mEPSCs between *Bbs4*^−/−^ and *Bbs4*^+/+^ mice. (N= 6; P = 0.24, Kolmogorov-Smirnov test)

### IGF-1R downstream signalling is dysregulated in *Bbs4*^−/−^ synaptosomes

It has been established that a number of tyrosine kinase receptors (RTK), including IGF, RET, TrkB, PDGF and EphB enhance dendritic growth and promote formation and maintenance of dendritic spines (7, 21). To assess the signalling of RTK in the brain of *Bbs4^−/−^* and *Bbs4^+/+^* mice we quantified the phosphorylation level of RTK receptors using a Phospho-RTK Array. Total brain extracts of P7 *Bbs4^−/−^* and *Bbs4^+/+^* mice were incubated with the membrane containing immobilised RTK antibody followed by detection of RTK phosphorylation by a pan anti-phospho-tyrosine antibody. Interestingly, phosphorylation levels of a number of RTKs were altered including insulin and IGF1 receptors (Fig.4a). Annotation and the summary of phosphorylation levels of all analysed RTK are given in Supplementary figure 5. We continued this study with the focus on IGF-1R/Insulin receptor (IR) signalling as it is known to have a profound effect on neuroplasticity in the CNS (7–9). Pull down experiments confirmed that phosphorylation of IGF-IR/InsulinR was decreased in P7 enriched synaptosomal fraction of *Bbs4^−/−^* mice (Fig.4b). Additionally, phosphorylation levels of Akt, a downstream target of canonical IGF signalling, were significantly reduced (Fig.4b). Next we tested phosphorylation level of IRSp58 protein, an adaptor protein that is phosphorylated by IR and IGF-1R (22) and is highly enriched in the postsynaptic density (PSD) of glutamatergic synapses (23). We found that phosphorylation of IRSp58 was significantly reduced in synaptosomal fraction of P7 *Bbs4^−/−^* mice (Fig.4b). Furthermore, as the activities of IGF-1R and IRSp58 depend on interaction with Rho family GTPases (24, 25), we investigated the activities of Rac1 and RhoA GTPases. We observed that activity of RhoA was increased and, concurrently, Rac1 activity was decreased in the enriched synaptosomal fraction of P7 *Bbs4^−/−^* mice (Fig.4c, d). We next assessed whether dysregulation of IGF-1 signalling in Bbs^−/−^ mice affects the levels of NMDA and AMPA receptors in the total and synaptosomal fractions of *Bbs4*^−/−^ and *Bbs4*^+/+^ mice by western blotting. We observed a significant increase in the level of NMDA and AMPA receptors in synaptosomal fraction of P7 *Bbs4^−/−^* mice, whereas the changes in the receptors level in the total brain fraction were not detected (Fig.4e). These data are consistent with our previous findings of increased mEPSC amplitudes in *Bbs4^−/−^* neurons (Fig.3e).

**Fig.4.**
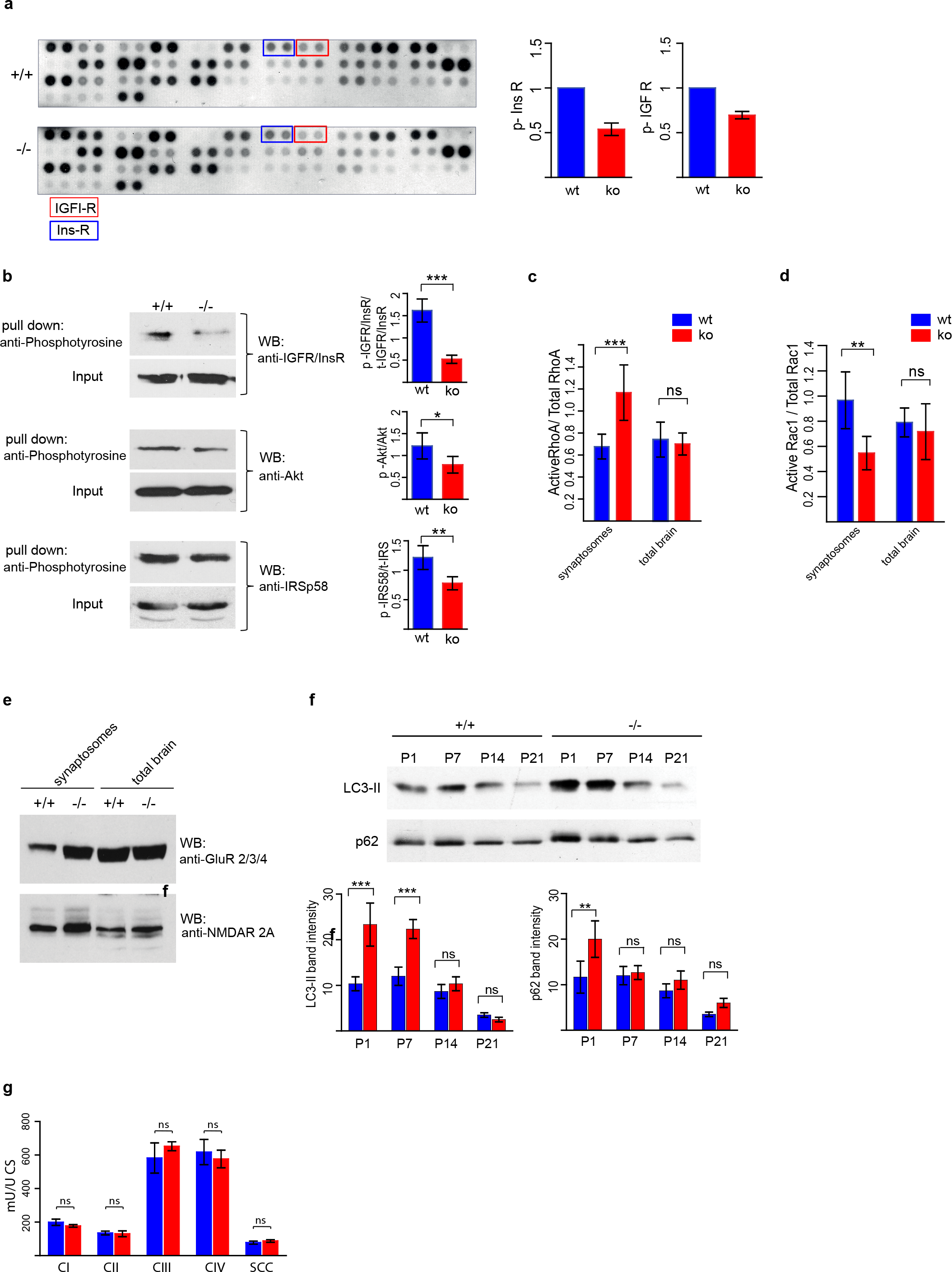
InsulinR/IGF1-R signalling is dysregulated in synaptosomal fraction of *Bbs4* knockout mice. **(a)** Phospho-RTK array reveals significant decrease in phosphorylation of Insulin and IGF1 receptors in P7 of *Bbs4*^−/−^ total brain protein extract (N=3, mean ± SD) **(b)** Pull down analysis shows aberrant IGF-1R downstream signalling. *Bbs4*^−/−^ and *Bbs4*^+/+^ enriched synaptosomal fractions were incubated with mouse anti-phosphotyrosine antibody overnight, followed by incubation with Dynabeads M-280 for 2 hours. Immunoblotting analysis of the proteins eluted from the beads was performed using anti-IGFR/InsR, anti-Akt, anti IRSp58 antibodies. Input: the total brain protein fraction before the incubation with anti-phosphotyrosine antibody, indicates the total level of IGFR/InsR, Akt and IRSp58 in *Bbs4*^−/−^ and *Bbs4*^+/+^ mice (N=3, mean ± SD) **(c, d)** RhoA and Rac1 G-LISA Activation Assays. Levels of activated RhoA **(c)** and Rac1 **(d)** were measured in the total brain extract and enriched synaptosomal fraction of P7 *Bbs4*^−/−^ and *Bbs4*^+/+^ mice (N=3, Mean ± SD; unpaired *t*-test) **(e)** Representative image of western blot analysis of NMDA and AMPA receptors levels in the total brain extract and enriched synaptosomal fraction of P7 *Bbs4*^−/−^ and *Bbs4*^+/+^ mice **(f)** Representative western blots of autophagy markers LC3-II and p62 in the synaptosomal fractions of *Bbs4*^−/−^ and *Bbs4*^+/+^ mice at P1, P7, P14 and P21 (N=3, mean ± SD). LC3-II and p62 levels were quantified by measuring Western blot band intensities using Image J programme (N=3, mean ± SD, unpaired *t*-test). Housekeeping genes (actin, GAPDH, etc) could not be used as normalization controls due to the changes in their gene expression levels in *Bbs4*^−/−^ mice. It was unexpected to see p62 increase in P1 synaptosomes in our experiment given the widely recognised notion that the level of p62 correlates inversely with autophagy, however, it is in line with the reports that p62 levels can be up-regulated during high autophagic flux due to a p62 protein multifunctional role (26) **(g)** Measurement of oxidative phosphorylation (OXPHOS) complex activities in the whole brain homogenates of *Bbs4*^−/−^ and *Bbs4*^+/+^ mice. CS, citrate synthase, CI, CII, CIII; mitochondria complex I, II, III; SCC, succinate:cytochrome c oxidoreductase (=complex II+ III combined); Units: mU:U CS; Raw data were normalised to citrate synthase; N=3, Mean ± SD, ns: not significant; unpaired *t*-test

One of the possible mechanisms of dendritic spine pruning is macroautophagy (*18*) process that is tightly regulated by IGF-1R signalling and small GTPases. To test whether autophagy is dysregulated in our *Bbs* model, we analysed the level of autophagy markers LC3-II and p62 in the enriched synaptosomal fractions isolated from *Bbs4^−/−^* and *Bbs4^+/+^* mice brains at different postnatal stages. We observed a significant increase in LC3-II level at P1 and P7 of *Bbs4^−/−^* mice (Fig.4f). Given the widely recognised notion that the level of p62 correlates inversely with autophagy, it was unexpected to see p62 increase in P1 synaptosomes in our experiment. However, it is in line with the reports that p62 levels can be up-regulated during high autophagic flux due to a p62 protein multifunctional role (26). To exclude mitochondrial dysfunction and oxidative stress as triggers of autophagic induction (27) we assessed the activities of respiratory chain complexes I, II, III, IV and SCC (complex II and III combined) in total brain homogenates of P7 *Bbs4^−/−^* and *Bbs4^+/+^* mice. Oxidative phosphorylation (OXPHOS) complex activities were determined and the results were normalized to the activity of citrate synthase (CS). We have found no significant differences in the activities of OXPHOS between *Bbs4^−/−^* and *Bbs4^+/+^* mice (Fig.4g) thus ruling out mitochondrial dysfunction as a cause of autophagy in BBS.

Our finding that aberrant IGF-1 signalling leads to the dysregulation of various cellular pathways that tightly control actin re-arrangements provide a possible explanation for spine density reduction in the absence of functional Bbs proteins.

### Synaptic localization of BBS proteins

To our knowledge, all previous studies confined the function of BBS proteins to structural and functional regulation of cilia. However, a reduction in dendritic spine density along with aberrant synaptic IGF receptor signalling and altered neurotransmitter receptor levels (NMDA and AMPA) suggested a potential synaptic role of Bbs proteins. Re-evaluation of our earlier mass spectrometric analyses of synaptosomal (28, 29) and crude synaptosomal fractions (30) of the rat cortex, dorsal striatum and dentate gyrus revealed the presence of Bbs1, Bbs2, Bbs4, Bbs5, Bbs7 and Bbs10 proteins in these fractions (Supplementary Table1).

To elaborate synaptic localization of BBS proteins biochemically, we enriched the cytosolic, detergent-soluble synaptosomal (DSS, pre-synapse enriched) and PSD fractions of synaptosomal preparations from the adult rat hippocampi using previously described method (31). Experimental procedure is summarised in the Supplementary Figure 6. Label-free MS1 intensity based LCMS quantitation revealed a high abundance of Bbs1, Bbs2, Bbs5, Bbs9 proteins in PSD fraction whereas Bbs7 was present mostly in the cytosolic fraction (Fig.5a, b). The low abundance Bbs4 protein was also unambiguously identified in the PSD fractions (Fig.5a,b). Immunofluorescence analysis of Bbs4 and Bbs5 protein localization confirmed the presence of Bbs punctae throughout the entire dendritic tree of mouse dissociated hippocampal neurons (Fig.5c and Supplementary Fig.7). Collectively, these data clearly indicate the presence of Bbs proteins in the neuronal processes and postsynaptic densities.

**Fig.5.**
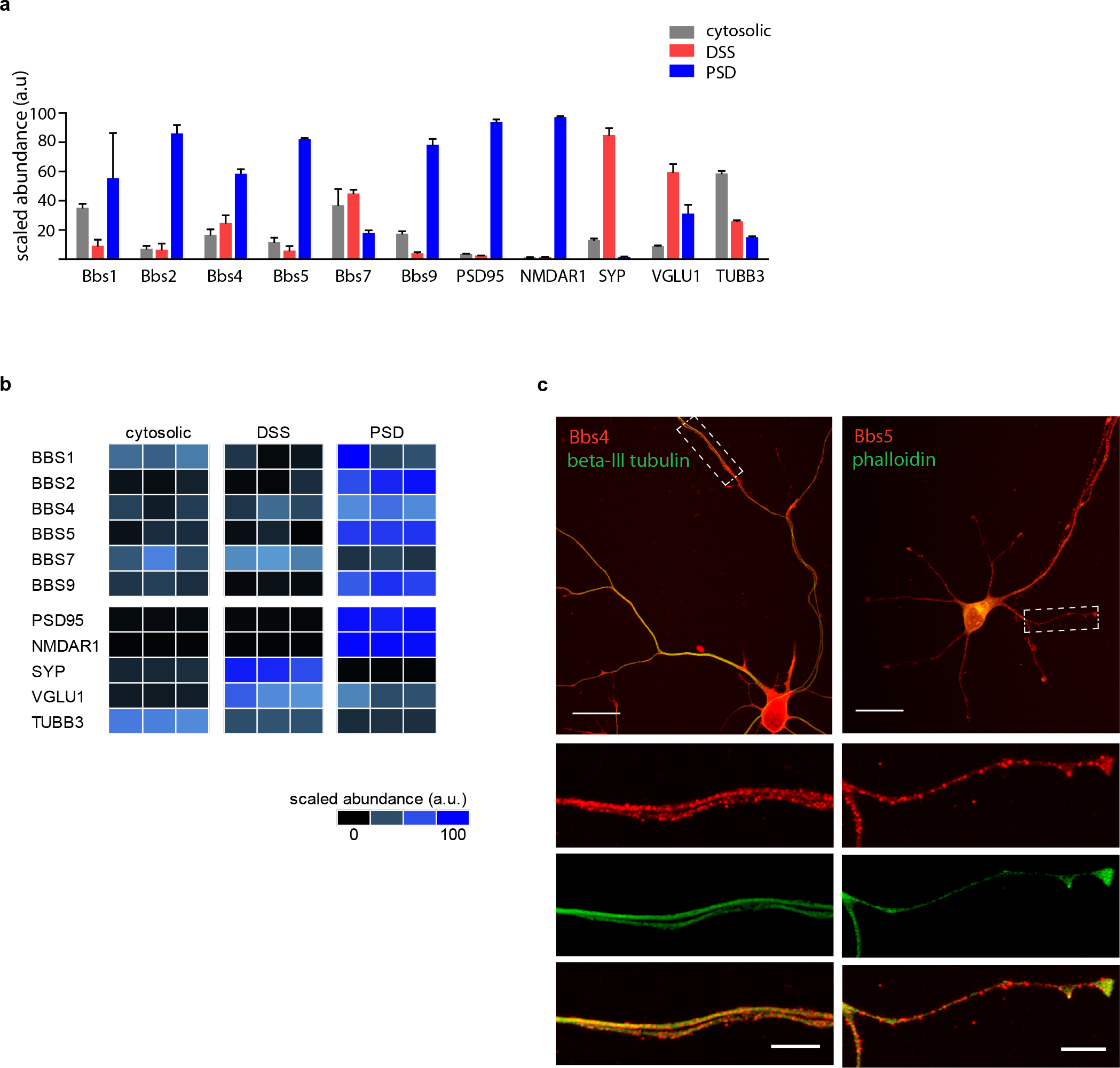
Biochemical and immunocytochemical profiling of BBS proteins. **(a)** Proteomic profile of the BBS proteins in biochemical fractions using nano-LC-MS/MS analysis. Protein levels of representative proteins for the following biochemical fractions of the rat hippocampus: cytosolic, detergent-soluble synaptosomal preparation (DSS, pre-synapse enriched), postsynaptic density preparation (PSD). Protein levels of representative synaptic markers were estimated from label-free LCMS analyses. Protein levels ofthe presynaptic (VGLU1, SYP) and postsynaptic (NMDAR1, PSD95) protein markers are enriched in DSS, and PSD preparations, respectively. The protein abundances illustrated in the heatmap are obtained from the total MS1 peptide intensities scaled to the mean of all the samples **(b)** Representative image of immunolabelling of Bbs proteins. Cultured mouse hippocampal neurons were immuno-labelled with Bbs4 and Bbs5 antibodies (red), phalloidin (green) and beta-III tubulin (green) after 6 days *in vitro* (DIV6). Scale bar 20 μm (top panel) and 5 μm (bottom panels).

### Voluntary exercise partly reverses dendritic spine loss in *Bbs4*^−/−^ mice

Aerobic exercise is known to promote spinogenesis and, even though the exact mechanisms are unclear, a number of studies suggested that it might work through several signalling molecules including IGF-1 and BDNF (32, 33). In the light of our finding of aberrant IGF-1 signalling we investigated whether voluntary wheel exercise is able to increase spinogenesis in *Bbs4^−/−^* mice. We analysed the spine density of DG neurons in 6 weeks old *Bbs4^+/+^*, *Bbs4^−/−^* and *Bbs4^−/−^* mice after 2 weeks of wheel exercise. We found that remarkably only 2 weeks of wheel exercise significantly increased the total spine density of *Bbs4^−/−^* mice DG neurons (Fig.6a,b). Analysis of spine density per branch order revealed increased density of spines in the 1^st^, 5^th^ and 6^th^ branch orders. Notably, although statistically not significant, there was a trend in spine density increase in the other branch orders (Fig.6c).

**Fig.6.**
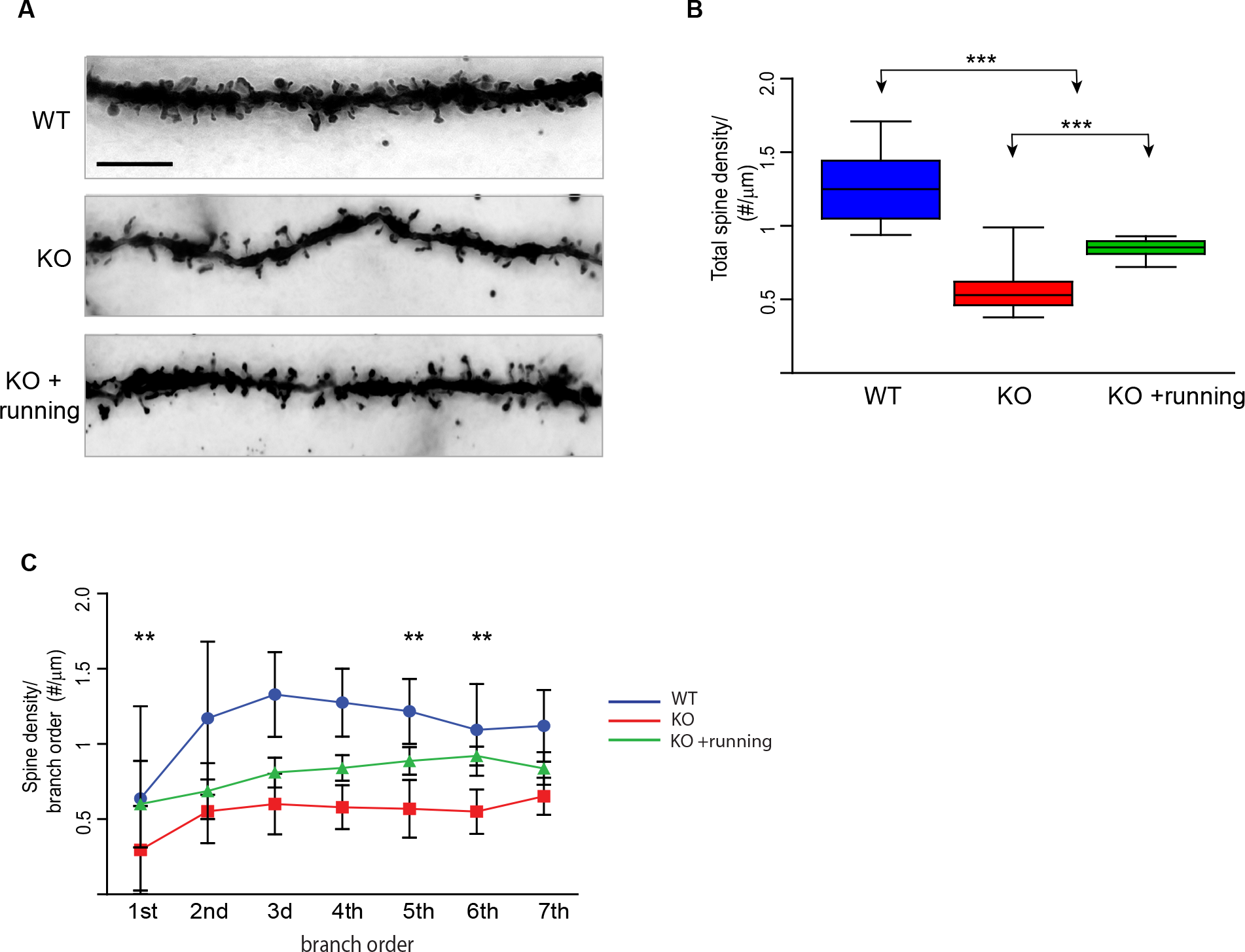
Two weeks of voluntarily wheel exercise increased dendritic spine density of DG hippocampal neurons in *Bbs4* knockout mice. **(a)** Representative images of Golgi-impregnated dentate granule (DG) of *Bbs^+/+^*, *Bbs4^−/−^,* and *Bbs4^−/−^* mice after 2 weeks of wheel-based exercise. (100x; scale bar 5 μm) **(b)** Total spine density of DG neurons of *Bbs^+/+^*, *Bbs4^−/−^,* and *Bbs4^−/−^* mice after 2 weeks of wheel-based exercise **(c)** Spine density per branch order of DG neurons of *Bbs^+/+^*, *Bbs4^−/−^,* and *Bbs4^−/−^* mice after 2 weeks of wheel-based exercise Biological samples: N_WT_=5; N_KO_=7; N_KO+running_=5. Total number of analysed cells: N_WT_=25; N_KO_=35, N_KO+running_=25. Mean ± SD, *P< 0.05. Unpaired *t*-test and one-way *ANOVA*, Tukey *post hoc* test

## Discussion

Our study reveals a previously unknown role for BBS proteins in neuronal function. We found significant morphological changes in dendritic spines and dendritic length in various brain regions of *Bbs* mouse models. Several studies have shown an association of neurodevelopmental and neuro-psychiatric disorders with morphological and physiological alterations in dendritic spines (2, 3) which prompted us to speculate that cognitive deficits in BBS patients are likely caused by the loss of dendritic spines. Our data provide several associative links between perturbed spine integrity and cognitive deficits in BBS: first, we show that *Bbs4^−/−^* mice display reduced contextual and cued fear memory. Secondly, loss of spines occurs in a spine sub-type-dependent manner, affecting mostly “thin” spines, which are thought to be specifically involved in learning processes, but not in memory maintenance (34). This finding is in good agreement with our clinical observation that learning difficulties in BBS patients are more prevalent than memory deficits (unpublished data). Finally, we found that the functional loss of different Bbs proteins appears to affect spines to different degree, for example, the *Bbs1 M390R* missense mutation has only a marginal effect on the spine loss. Together, these data support the idea that dendritic spine aberration might be an essential contributing factor to cognitive deficits in BBS.

Loss of dendritic spines may be attributed to a number of mechanisms including reduced synaptic activity, mitochondrial dysfunction etc. Our data indicate that as the intrinsic properties of Bbs4^−/−^ neurons were unaffected and mEPSC amplitudes were significantly larger in Bbs4^−/−^ neurons, it is it unlikely that reduced neuronal activity might be the cause of reduced spine density suggesting compensatory responses to the spine loss. Similarly, we did not obtain any mitochondrial dysfunction in *Bbs4^−/−^* brain. Strikingly, we found that several tyrosine kinase receptors signalling was affected including IGF-1R and ephrinB2 receptor. A number of studies reported the effect of IGF-1R and ephrin B2 receptor signalling on dendritic growth, formation, maintenance, and stabilization of dendritic spines (7, 35). Our present work shows that disruption in IGF-1 signalling initiates a cascade of downstream events such as increased autophagy, reduced phosphorylation of IRSp58 and aberrant activity of small GTPases, Rac1 and RhoA. All these events are known to have a diminishing effect on the spine density. Thus, our finding of aberrant cellular pathways that tightly control actin re-arrangements provide a plausible explanation for spine density reduction in the absence of functional Bbs proteins.

How ciliary Bbs proteins affect the signalling of the receptors integrated into spine membrane? Our study revealing the presence of BBS proteins in post synaptic density of the spines. Furthermore, comparison between primary cilia and dendritic spines highlights remarkable parallels between protein composition, membrane domain architecture and dynamic nature of primary cilia and dendritic spine assembly/ disassembly (36). Collectively, it is suggestive that BBS proteins might have similar function in both structures. Therefore, we propose a model in which BBS proteins are localised to the dendritic spines and stabilize microtubule polymerization (invasions) into dendritic spines thus facilitating the transport of the signalling receptors to the spine membrane (Fig.7). It would require further investigations to determine whether BBSome complex is present in the spines similar to primary cilia.

**Fig.7.**
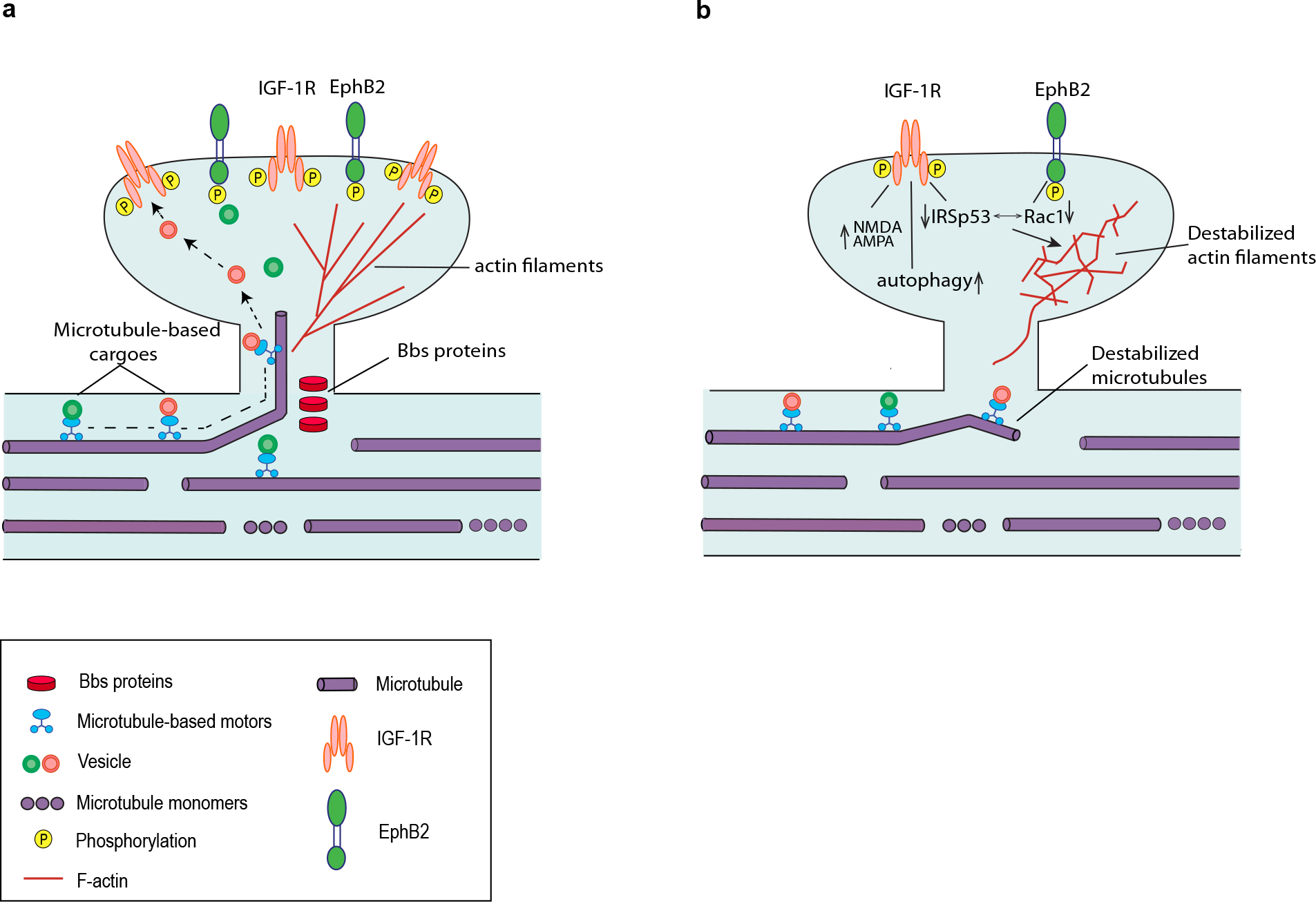
Model of the role of Bbs proteins in synapses. This simplified schematic is based on the data shown in this paper and on the previously described role of Bbs proteins in primary cilium. It illustrates possible mechanisms by which Bbs proteins might impact synapse plasticity **(a)** In the presence of Bbs proteins at the dendritic shafts microtubules are anchored facilitating receptor transport to the plasma membrane of the spines. Bbs proteins might be also involved in the sorting of the receptors to be transported to the spine membrane **(b)** In the absence of Bbs proteins the microtubule anchoring and receptor sorting is destabilized leading to reduced receptor trafficking to the spine membrane. The receptor transport to the spines becomes dependent on the other transport mechanisms (e.g. exocytosis of vesicles in the dendritic shafts or a myosin-based transport (40). As a result, reduced receptor signalling (e.g. IGF-1R) triggers a number of events leading to destabilization of actin filaments and eventually dendritic spines

Finally, we show that that in *Bbs4* mice model spine density can be increased by only two weeks of voluntarily wheel exercise. Accumulating evidences suggest that voluntary wheel running has an enhancing effect on the dendritic spine density and can even reverse spine loss in the mouse model of Parkinson’s disease (37, 38). Although the exact mechanisms of neuroplasticity enhancement during aerobic exercise remains unclear, several important signalling molecules including IGF-1 and BDNF were shown to be implicated (32, 33). In the light of this evidence, it is interesting that Bbs4 protein was found to be necessary for TrkB receptor localization to the cilia membrane and activation by BDNF(15). Considering structural similarity between primary cilia and dendritic spines it is plausible that Bbs4 proteins are involved in the localization of TrkB receptors on the spine membrane.

Even though it was not investigated in this work, our data may be pivotal in understanding already known interaction between BBS proteins and the Disrupted-in-schizophrenia 1 (DISC1) protein, the disruption of which can result in a wide a range of psychiatric problems including schizophrenia, bipolar disorder and major depression (39). As loss of spines is relevant to many brain disorders including neurodegeneration, probing the synaptic role of BBS proteins will contribute to a deeper understanding of the aetiology of these disorders.

## Supporting information

Bbs4 reconstructed neurons

## Acknowledgements

We thank International Mouse Phenotyping Consortium at Harwell for providing C57BL/6NTac-Bbs5^tm1b(EUCOMM)Wtsi^/H strain model and for assistance in experimental techniques. This study was supported by grants from Biotechnology and Biological Sciences Research Council (BBSRC) (BB/M020991/1) and New Life Foundation (14-15/20), UK.

## Authors’ contributions

N.H. performed the experiments and generated the figures. C.S-H. performed the electrophysiology experiments and assisted with manuscript preparation. F.S. carried out proteomics experiments, analysed the data and prepared proteomics figures. L.C. isolated mouse hippocampal neurons and performed immunocytochemical experiments. D.M performed multiphoton microscope imaging. M.S., L.B., S.W., and P.N performed behavioural tests. R.R performed mitochondria function tests. E.F. assisted with manuscript preparation. M.C.W guided dendritic spine analysis and made a video of reconstructed neurons. P.S. guided in vitro experiments. G.L. guided proteomics experiments. M.H. guided electrophysiology experiments and assisted with manuscript preparation. P.L.B assisted with manuscript preparation. S.C-S conceived the study, performed experiments, generated the figures, and wrote the manuscript.

Correspondence and requests for materials should be addressed to S.C-S

## Methods

### Mice and ethics statement

Animal maintenance, husbandry, and procedures are defined and controlled by the Animal (Scientific Procedures) Act 1986. All animal experimentations has been carried out under licences granted by the Home Secretary (PIL No. 70/7892) in compliance with Biological Services Management Group and the Biological Services Ethical Committee, UCL, London, UK. Mice were group housed in IVC cages and were kept on a 12-h light-dark cycle with ad libitum access to food and water.

*Bbs1* M390R knock in model was purchased from Jackson Laboratory, *Bbs4* gene trap model we received as a part of previous collaborative work (41), and targeted knockout C57BL/6NTac-Bbs5^tm1b(EUCOMM)Wtsi^/H strain model was received from MRC Harwell as part of the International Mouse Phenotyping Consortium. *Bbs1* and *Bbs4* mice were backcrossed with C57BL/6Ntac strain for 5 generations to keep the background consistent with *Bbs5* mice obtained from MRC Harwell.

### Statistical analysis

Analysis with two groups were performed using an unpaired, two-tailed Student’s *t*-test. Analysis with more than two groups and with one variable were performed using one-way analysis of variance (*ANOVA*) and Tukey’s *post hoc* tests. Kolmogorov-Smirnov test was used to determine the cumulative distribution function of a continuous random variables such as frequency and amplitudes of mEPSCs.

### Data Availability

The authors confirm that the data supporting the findings of this study are available within the article and its Supplementary material. Raw data can be requested from the corresponding author.

## Supplementary Information

### Methods

#### Dendritic spine analysis

The brains were stained by commercially available Golgi-Cox impregnation kit (NeuroDigiTech). The coronal sections were prepared that covered the anterior-to-posterior axis of the brain and the regions of interest (ROI) (DG, BLA, and LV pyramidal neurons) were chosen and analysed using a stereology-based software, called NeuroLucida, v10 (Microbrightfield, VT), installed on a Dell PC workstation that controlled Zeiss Axioplan 2 image microscope with Optronics MicroFire CCD camera (1600 x 1200), motorized X, Y, and Z-focus for high-resolution image acquisition and digital quantitation. Five or six neurons per ROI were chosen for the analysis. The sampling process was conducted as follows: The investigators i) previewed the entire rostro-caudal axis of ROI, with low-magnification Zeiss objectives (10x and 20x), ii), compared and located those with the least truncations of distal dendrites as possible under high-magnification Zeiss objectives (40x and 63x), iii), and then iv) used a Zeiss 100x objective with immersion oil to perform 3D dendritic reconstruction, followed by continuous counting of spines throughout the entire dendritic trees. The criteria for selecting candidate neurons for analysis were based on (i) visualization of a completely filled soma with no overlap of neighbouring soma and completely filled dendrites, (ii) the tapering of most distal dendrites; (iii) the visualization of the complete 3-D profile of dendritic trees using the 3-D display of imaging software. Neurons with incomplete impregnation and/or neurons with truncations due to the plane of sectioning were not collected. Moreover, cells with dendrites labelled retrogradely by impregnation in the surrounding neuropil were excluded. With the systematic registration and digital monitoring, the software was able to accurately record every step of the tracing/contouring and generate a 3-D reconstructed dendritic morphology for subsequent spine counting. Automatic navigation of the digital probes with registered x-y-z coordinates of each 2-D image stack enabled creating a complete 3-D neural reconstruction for the dendrograms, spine density and Sholl analysis. It is noted that only spines orthogonal to the dendritic shaft were readily resolved and included in this analysis, whereas spines protruding above or beneath the dendritic shaft were not sampled (see below). This principle was remained consistent throughout the course of analysis. Also, due to inevitable truncations and shrinkage after impregnation process and optical limitation to resolving most distal dendrites in deep z-axis, under-estimates of the actual dendritic lengths and spine numbers would be expected. The above limitations, however, do not affect the comparison of morphological properties between animals.

#### Mass spectrometric analysis

Hippocampi from fresh Sprague-Dawley rat brains (n = 4) were dissected and stored at −80°C. Crude synaptosomes were prepared from tissues according to previously described protocols (42). Biochemical fractionation of crude synaptosomes into cytosolic, detergent-soluble (pre-synapse enriched) and post-synapse-enriched synaptosomal were prepared as previously described (31). Methods for sample preparation for liquid chromatography followed by tandem mass spectrometry (LC-MS/MS) were implemented as previously described (28). Peptide analyses were performed using a single-shot LC-MS/MS approach with a 4-hr gradient using a Dionex Ultimate 3000 system (Thermo Fisher Scientific) coupled to a Q-Exactive Plus mass spectrometer (Thermo Fisher Scientific, Germany) with LCMS parameters as described previously (43). All MS-MS2 spectra were searched against UniProtKB/Swiss-Prot rat protein database version v 2016.04.14. Parameters for protein identification and label-free quantification workflow based on the Minora algorithm through Proteome Discoverer 2.2 platform as previously described (43). Abundances are scaled according to the mean abundance of the samples.

#### Electrophysiology

Transverse 300μm-thick slices were cut from the hippocampi of 3 week-old mice with a VT1200 vibratome (Leica Microsystems, Wetzlar, Germany). The animals were anaesthetized with isoflurane added to the inspiration airflow (4–5%; Abbott, Ludwigshafen, Germany) and killed by decapitation, in accordance with national and institutional guidelines. The cutting solution was sucrose-based containing (in mM) 87 NaCl, 25 NaHCO_3_, 2.5 KCl, 1.25 NaH_2_PO_4_, 75 sucrose, 0.5 CaCl_2_, 7 MgCl_2_ and 10 glucose (equilibrated with 95% O_2_ –5% CO_2_). After cutting, slices were kept at 35°C for 30 min and then stored at room temperature in artificial cerebrospinal fluid (ACSF) containing (in mM): 125 NaCl, 25 NaHCO_3_, 2.5 KCl, 1.25 NaH_2_PO_4_, 2 CaCl_2_, 1 MgCl_2_, 25 glucose (equilibrated with 95% O_2_–5% CO_2_).

For electrophysiological experiments, slices were continuously superfused with ACSF. Patch pipettes (4–8 MΩ) were pulled from borosilicate glass tubing with 1.5 mm outer diameter and 0.86 mm inner diameter (Warner, Hamden, USA). The pipettes were filled with a solution containing 130 mM potassium gluconate, 20 mM KCl, 2 mM MgCl_2_, 4 mM K_2_ATP, 0.3 mM NaGTP, 10 mM sodium phosphocreatine and 10 mM Hepes. The pH was adjusted to 7.3 by adding KOH. Voltage signals and currents were measured with a Multiclamp 700B amplifier (Molecular Devices, Palo Alto, CA, USA), filtered at 5 kHz and digitized at 20kHz using a Digidata 1550 interface (Molecular Devices, Palo Alto, CA, USA). Data analysis was performed with Stimfit (Guzman et al., 2014). Membrane potentials not corrected for liquid junction potentials. All recordings were made at near-physiological temperatures (32–34°C). Spontaneous mEPSCs were recorded at a membrane potential of −80 mV in the presence of 0.5 μM TTX, 1 μM gabazine and 25 μM D-AP5. For detection and analysis of miniature synaptic events a template-matching algorithm was used, implemented in Stimfit (Jonas *et al.* 1993; Clements & Bekkers, 1997; Guzman et al., 2014).

#### Contextual fear conditioning test and cue fear conditioning

Context and cue-dependent fear conditioning experiments were performed using a fear conditioning chamber (bought from Ugo Basile). Mice were trained and tested in the chamber with clear plastic walls and ceiling and a standard grid floor. On Day 1 mice were placed into the fear conditioning chamber for 616.5 s. After 150 s a 5 s tone was played, followed immediately by a 0.5 s, 0.5 mA shock. The tone–shock pairing was repeated another two times at 150 s intervals. For the contextual conditioning test, on day 2, mice were placed in exactly the same arenas as Day 1 for 300 s. No tones or shocks were presented. Video tracking was recorded. For the cued conditioning test, on day two four hours after the contextual test, mice were placed in an altered context (round arenas, flat bottomed instead of a grid, stripy walls, added vanillin scent around the top, reduced lux) for 180 s with no tones or shocks. At 180 s a 5 s tone was added for a total of three tones with 60 s played 60 s intervals. Video tracking was recorded and scoring of freezing behaviour was automatically performed and analysed by Any-Maze software.

#### Phospho-RTK array, pull down experiments, and western blotting

Total brain extracts and enriched synaptosomal fractions from the whole brain were isolated from E19.5 embryos and P1, P7, P14 and P21 postnatal mice using RIPA buffer and Syn-PER™ Synaptic Protein Extraction Reagent (ThermoFisher Scientific) with protease inhibitors cocktail (ThermoFisher Scientific) and in accordance to standard procedure and manufacture’s protocol. Protein concentration was determined using Pierce BCA Protein Assay Reagent (ThermoFisher Scientific). Phospho-RTK array (R&D systems) was performed using total brain protein extracts of P7 *Bbs4^−/−^* and *Bbs4^+/+^* and in accordance to manufacture protocol. For pull-down experiments enriched synaptosomal fractions of *Bbs4^−/−^* and *Bbs4^+/+^* were incubated with mouse anti-phosphotyrosine (BD Transduction) antibodies overnight at 4°C, incubated with sheep anti mouse IgG Dynabeads M-280 (ThermoFisher Scientific) for 2h at 4°C and analysed by Western blotting using anti IRS58, anti-IGFR/InsR (Cell Signaling), anti-Akt (Cell Signaling), anti-IRS58 (Abcam) antibodies. The levels of AMPARs and NMDARs in total brain and enriched synaptosomal fractions were analysed using anti-GluR 2/3/4 (Cell Signaling), anti-NMDA subunit 2A (Life Technologies) antibodies and standard western blotting procedure. The intensities of the bands were analysed by ImageJ software.

The activities of Rac1 and RhoA small GTPases in the total brain and enriched synaptosomal fractions were measures using G-LISA RhoA activation assay (Cytoskeleton, BK124) and G-LISA Rac1 activation assay (Cytoskeleton, BK128). The activity of RhoA was normalised to the total level of RhoA (Cytoskeleton, BK150).

#### Immunofluorescence and image acquisition

Primary hippocampal cultures were prepared from embryonic day 18 C57BL/6Ntac mouse embryos and cultured at medium-density (100 cells per mm^2^) in N2/B27 medium for 6 days in vitro (DIV)(44). Dissociated neurons were fixed with 4% formaldehyde, permeabilized with cold 100% methanol, blocked with 1% horse serum and incubated with primary antibodies at RT for 1h. Primary antibodies against tubulin (Tuj-1) (Chemicon), Bbs5 and Bbs4 (ProteinTech), Alexa Fluor phalloidin 488 (Invitrogen) were used. Secondary antibodies Alexa 488, Alexa 568 were from Invitrogen. The images were taken using Zeiss LSM 880 upright confocal microscope with Airyscan, 63x/NA1.4 Plan Apo Oil lens.

#### Mitochondria Function

Mitochondrial respiratory chain enzyme and citrate synthase activities in 600g supernatants of whole brain homogenates (10% *w*/*V* in 0.25 M sucrose, 2 mM EDTA, 10 mM K_2_HPO_4_/KH_2_PO_4_, pH 7.4) were measured following previously described procedures (45) and references therein on a Konelab 20XT spectrophotometric analyzer (ThermoFisher Scientific).

---

**Supplementary Table 1**

Published and current proteomics data where BBS proteins were identified. LCMS-based proteomic analyses of rat synaptosomal and membrane preparations

**Supplementary Video 1**

**Reconstructed hippocampal *Bbs4*^−/−^ neurons show a significant reduction in total dendritic spines when compared to *Bbs*^+/+^ neurons.** For 3D neuron reconstruction (Neurolucida, MBF), the image stacks of representative neurons per group were selected in order to align serial contoured objects, including the soma, apical and basal dendrites and associated spines. The module “3D visualization” of Neuroludica software was turned on in order to automatically generate 3D visualization of representative neurons.

**Supplementary Fig.1**

**(a-e) Sholl analysis of DG, BLA and Layer V frontal cortex neurons of 6 weeks old *Bbs4* mice**. Frequency of intersections per 30 μm interval in DG **(a)**, apical dendrites of layer V pyramidal neurons **(b)** and basal dendrites of layer V pyramidal neurons **(c),** apical dendrites of BLA **(d)**, basal dendrites of BLA **(e)**. (N_WT_=5; N_KO_=7, Mean ± SD, *P< 0.05). One-way *ANOVA*, Tukey *post hoc* test

**(f-j)** Dendritic length of DG, BLA and layer V neurons. (Biological samples: N_WT_=5; N_KO_=7; Total number of analysed cells: N_WT_=25; N_KO_=35, for DG; Biological samples: N_WT_=3; N_KO_=3; Total number of analysed cells: N_WT_=15; N_KO_=15 for BLA and LV) Mean ± SD, *P< 0.05. Unpaired *t*-test

**Supplementary Fig.2**

**Defects in dendritic morphology of DG granule cells of 3 weeks old *Bbs5* knockout and *Bbs1* M390R knock in models**

**(a**) Representative images of Golgi-impregnated dentate gyrus (DG) granule cells of *Bbs5* and *Bbs1* M390R models.

**(b-f)** Analysis of DG granule cells of *Bbs5* mice. **(b)** total spine density, **(c)** dendritic length, **(d)** spine density per branch order, **(e)** spine density per 30 μm interval, **(f)** frequency of intersections per 30 μm interval

**(g-k)** Analysis of DG granule cells of *Bbs1*M390R knock-in mice. **(g)** total spine density, **(h)** dendritic length, **(i)** spine density per branch order, **(j)** spine density per 30μm interval, **(k)** frequency of intersections per 30 μm interval.

Biological samples: N_WT_=5; N_KO_=5; Total number of analysed cells: N_WT_=25; N_KO_=25, Mean ± SD, *P< 0.05, P***<0.01. One-way *ANOVA*, Tukey *post hoc* test. Scale bar 5 μm. # represents a significant level if un-paired, two-tailed *t* test is used (p<0.05).

**Supplementary Fig. 3**

**‘Thin’ spines are abandoned on DG neurons of 6 weeks old *Bbs4*knockout mice**

**(a)** Illustration of dendritic spine sub-classes. Adopted from Oliver von Bohlen und Halbach, Annals of Anatomy(18)

**(b)** Total count of ‘thin’, ‘stubby’, ‘mushroom’, ‘filopodia’ and ‘branched’ spines

**(c)** Total dendritic density of thin, stubby, mushroom, filopodia and branched spines. Density is calculated as the number of spines per micrometre of dendrite. Boxed panel shows the reduction in dendritic length.

**(d)** Sholl analysis of ‘thin’ and ‘mushroom’ spines (N_mice/WT_=3; N_mice/KO_=3, N_cells/WT_=15, N_cells/KO_=15, Mean ± SD, ***P<0.001; **P<0.01; *P< 0.05). One-way *ANOVA*, Tukey *post hoc* test

**Supplementary Fig. 4**

**Dendritic spine density is reduced in DG neurons of *Bbs4* knockout mice at P21 but not at E19.5**

**(a)** Representative images of Golgi-Cox impregnated dentate granule (DG) of *Bbs4* KO and WT mice at E19.5 and P21 (100x; scale bar 5 μm)

**(b-f)** Analysis of DG neuron morphology at E19.5. Total spine density **(b)**, dendritic length **(c)**, spine density per branch order **(d)**, spine density per 30 μm interval **(e)**, frequency of intersections per 30 μm interval **(f)**

**(g-k)** Analysis of DG neuron morphology at P21. Total spine density **(h)**, dendritic length **(i)**, spine density per branch order **(j)**, spine density per 30 μm interval **(k)**, frequency of intersections per 30 μm interval **(l)**

(N_mice/WT_=3; N_mice/KO_=3, N_cells/WT_=24, N_cells /KO_=24, Mean ± SD, ***P<0.001; **P<0.01; *P< 0.05). One-way *ANOVA*, Tukey *post hoc* test except for b, c, h, i where unpaired *t*-test was used

**Supplementary Fig.5**

**RTK phosphorylation assay of enriched synaptosomal fractions of P7 *Bbs4* knockout and control mice**

**(a)** Representative image of Phospho-RTK array. Blue and red squares indicate the position of Ins-R and IGF-1R, respectively.

**(b)** Quantitative dot-blot analysis reveals significant decrease in phosphorylation of a number of RTK in P7 of *Bbs4*^−/−^ mice brain(N=3, mean ± SD); unpaired *t*-test; ImageJ software.

**Supplementary Fig.6**

Experimental Workflow. Cytoplasmic and Synaptosomal fractions were isolated from the cortices of three Wistar rats. Synaptosomes were further separated into detergent soluble synaptosomal (DSS) and postsynaptic density (PSD) fractions. A FASP protocol adapted for synaptic membrane proteins is coupled to a gel-free LCMS to allow analyses of synaptosomal fractions. Database searching was performed with search engines against the rat SwissProt protein database. Quantitative information were determined using the software tools, Proteome Discoverer and Isobar.

**Supplementary Fig.7**

**Validation of the specificity of *Bbs4* and *Bbs5* antibodies**

**(a)** List of published validations of *Bbs4* (12766-1-AP) and *Bbs5* (14569-1-AP) ProteinTech antibodies used in this study

**(b)** Protein extracts from the brain of the wild-type and knockout *Bbs4* and *Bbs5* mice were immunoblotted with Bbs4 and Bbs5 antibodies as indicated. Approximate molecular weights are listed on the left side. Gapdh was used as the loading control. Wild type *Bbs4* and *Bbs5* mice showed a specific single band. Western blot with Bbs4 and Bbs5 antibodies did not detect any specific band in *Bbs4* and *Bbs5* knockout mice.

