## Supplementary figures for "Loss of Bardet-Biedl syndrome proteins causes synaptic aberrations in principal neurons"

**Supplementary Fig.1**

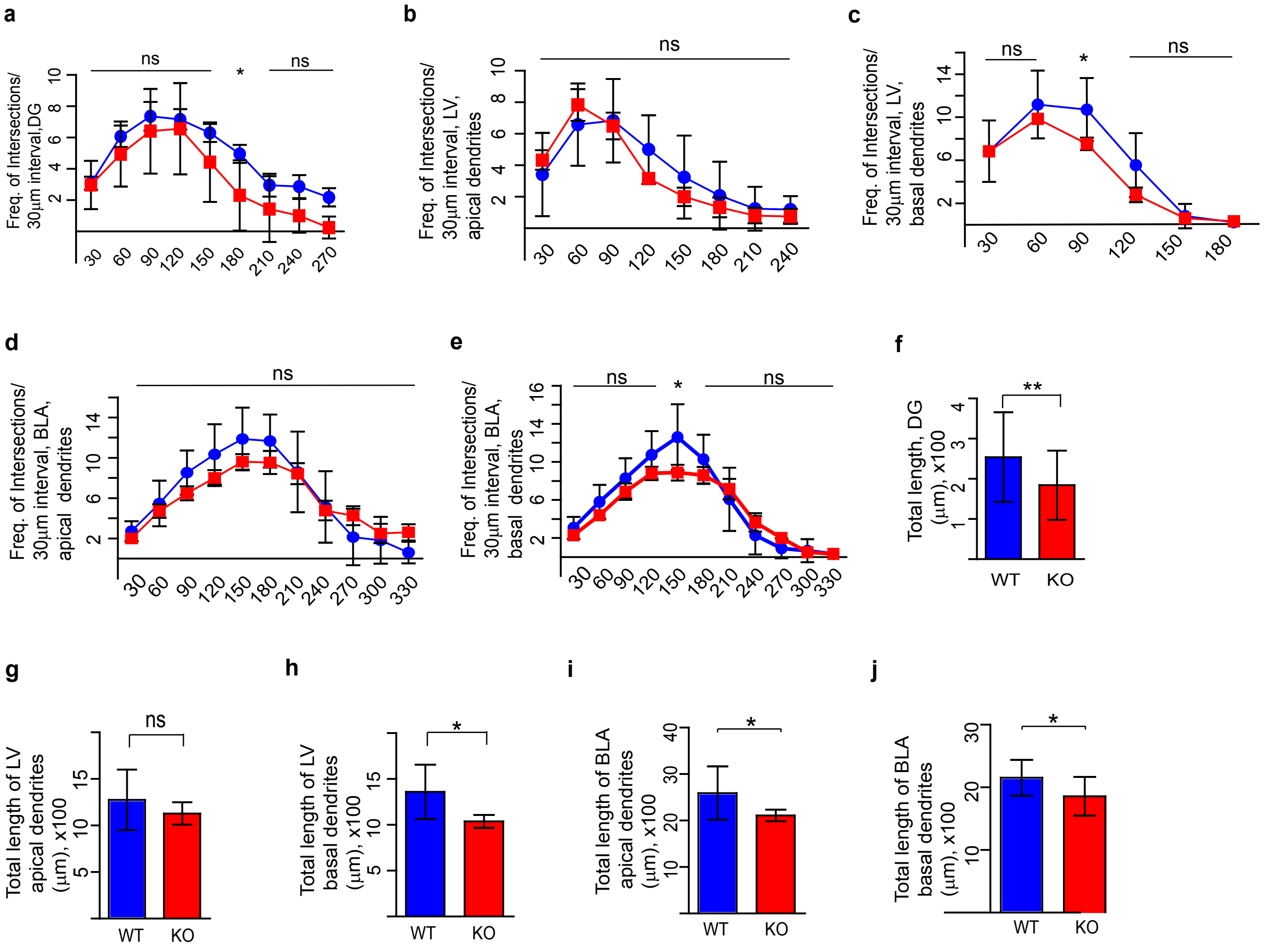

#### Supplementary Fig.2

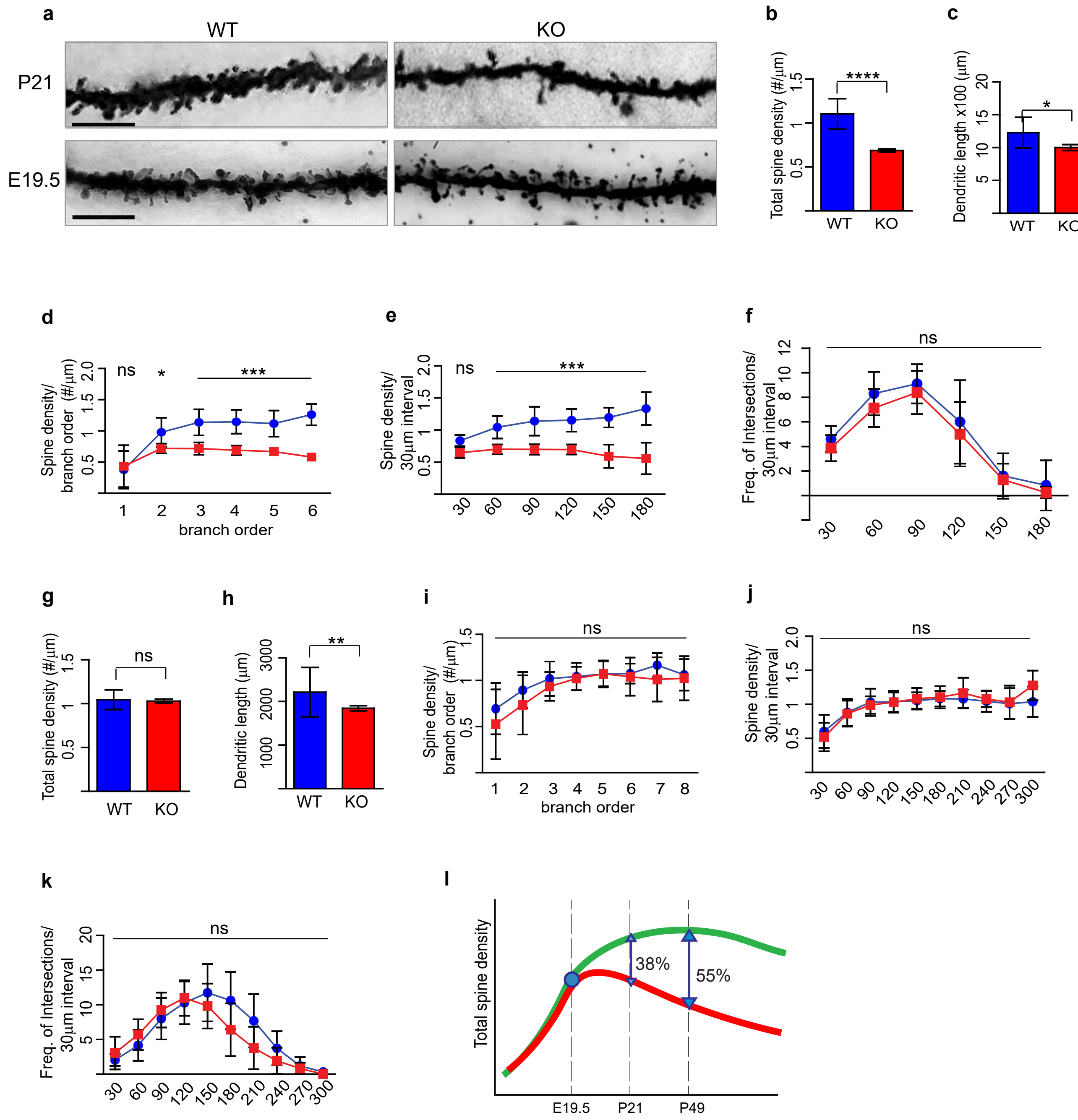

### Supplementary Fig.3

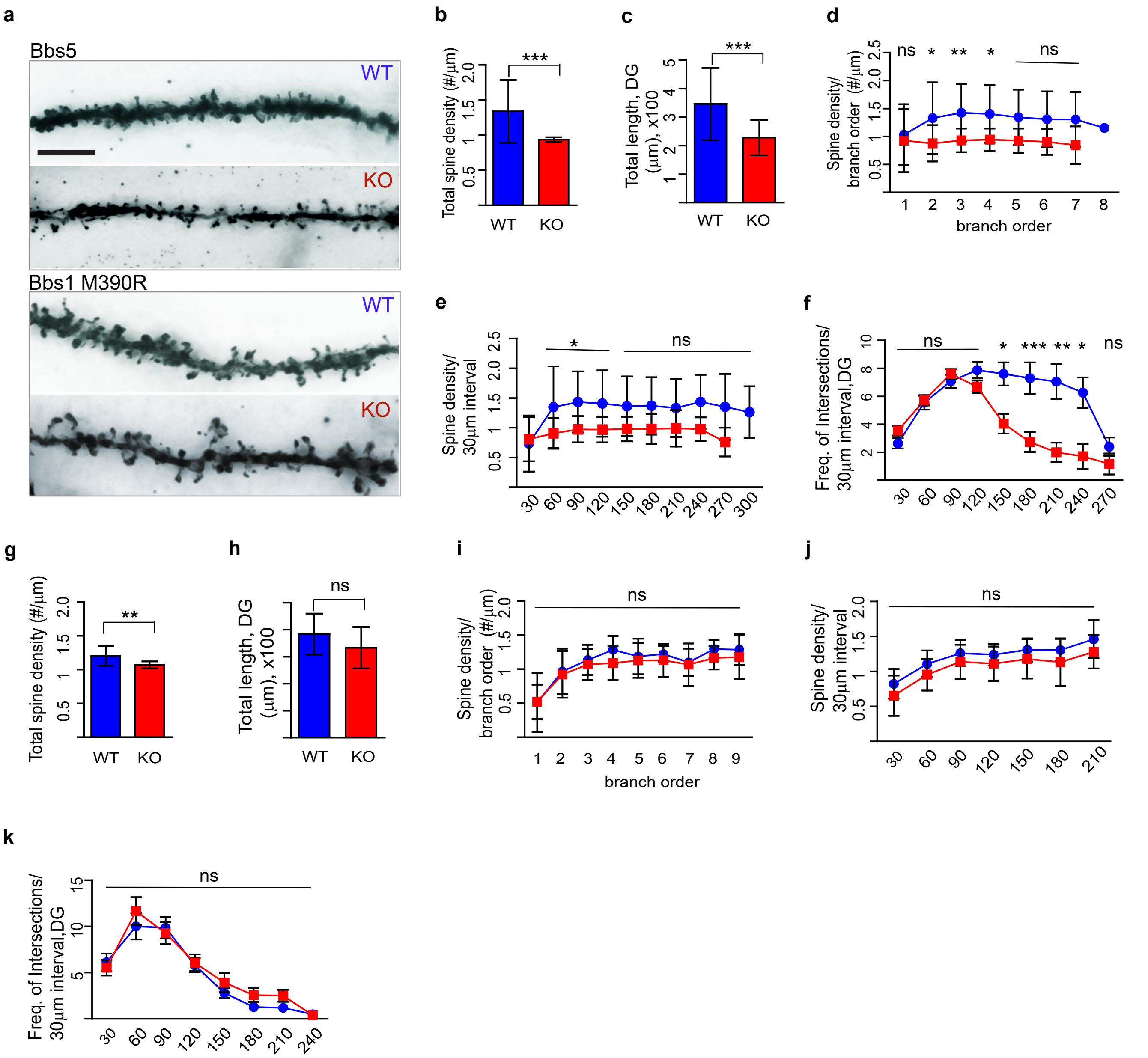

#### Supplementary Fig. 4

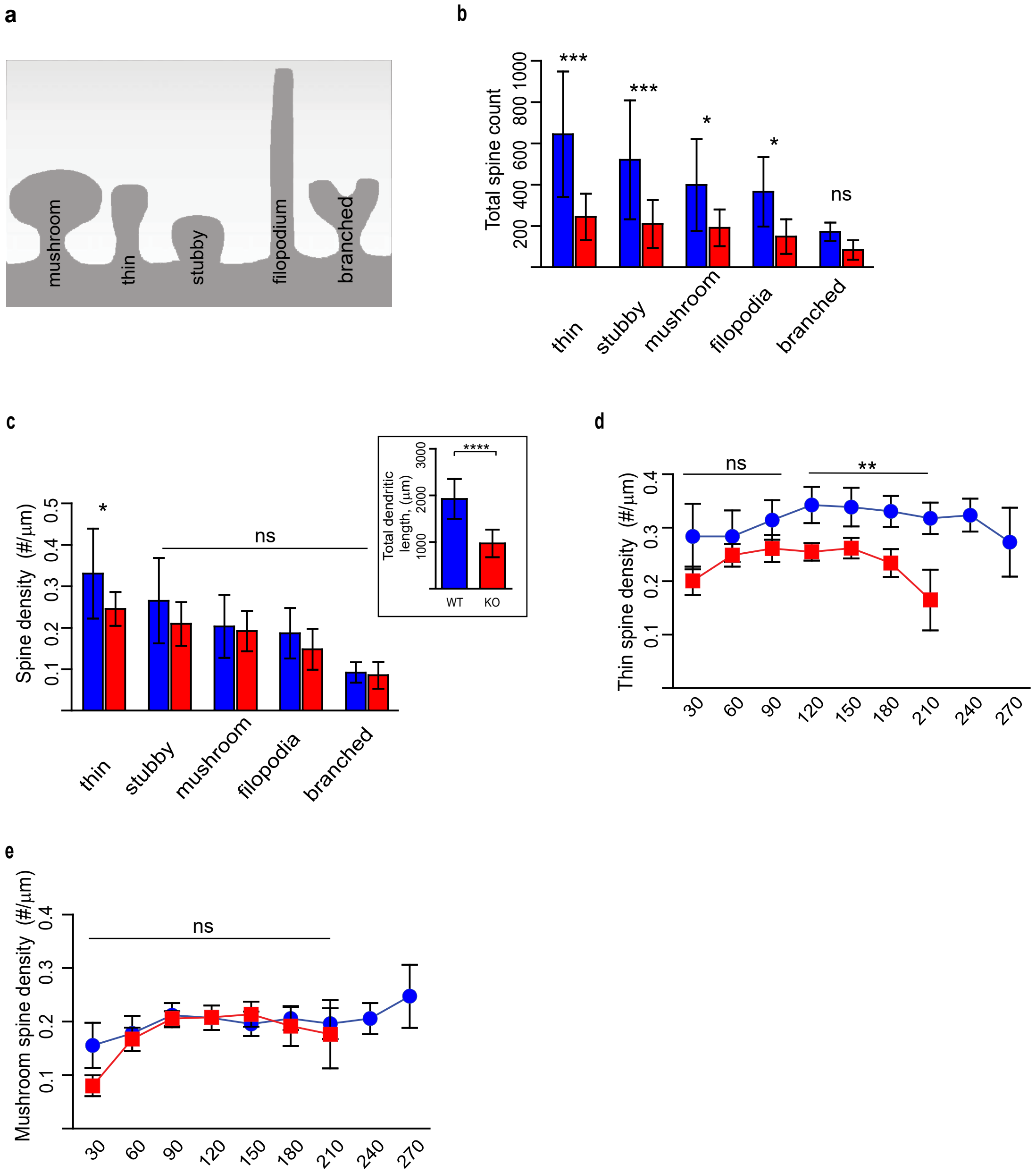

#### Supplementary Fig.5

**a**

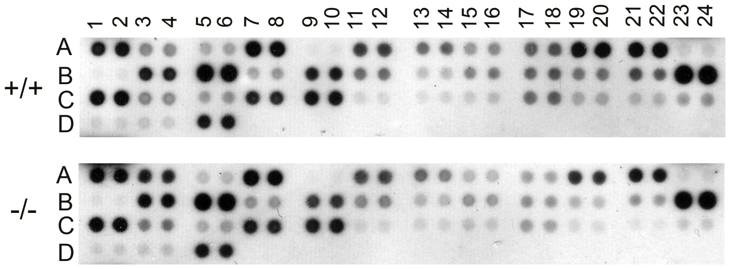

| | | | | | |
|---|---|---|---|---|---|
| **A1, A2** | EGF R | **B3, B4** | MSP R | **C5, C6** | VEGF R2 |
| **A3, A4** | ErbB2 | **B5, B6** | PDGF Rα | **C7, C8** | VEGF R3 |
| **A5, A6** | ErbB3 | **B7, B8** | PDGF Rβ | **C9, C10** | MuSK |
| **A7, A8** | ErbB4 | **B9, B10** | SCF R | **C11, C12** | EphA1 |
| **A9, A10** | FGF R2 | **B11, B12** | Flt-3 | **C13, C14** | EphA2 |
| **A11, A12** | FGF R3 | **B13, B14** | M-CSF R | **C15, C16** | EphA3 |
| **A13, A14** | FGF R4 | **B15, B16** | c-Ret | **C17, C18** | EphA6 |
| **A15, A16** | Insulin R | **B17, B18** | Tie-1 | **C19, C20** | EphA7 |
| **A17, A18** | IGF-IR | **B19, B20** | Tie-2 | **C21, C22** | EphA8 |
| **A19, A20** | Axl | **B21, B22** | TrkA | **C23, C24** | EphB1 |
| **A21, A22** | Dtk | **B23, B24** | TrkB | **D1, D2** | EphB2 |
| **A23, A24** | Mer | **C1, C2** | TrkC | **D3, D4** | EphB4 |
| **B1, B2** | HGF R | **C3, C4** | VEGF R1 | **D5, D5** | EphB6 |

**b**

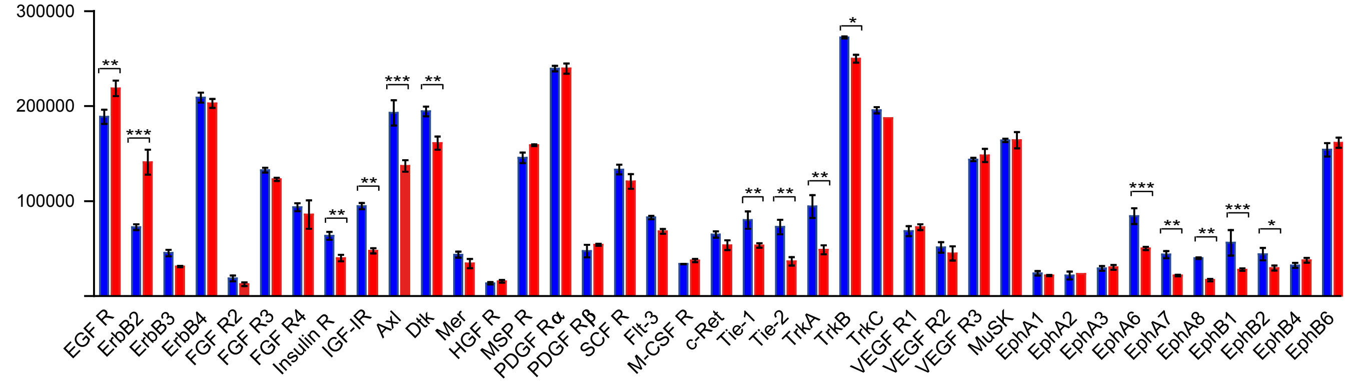

**Fig. 6**

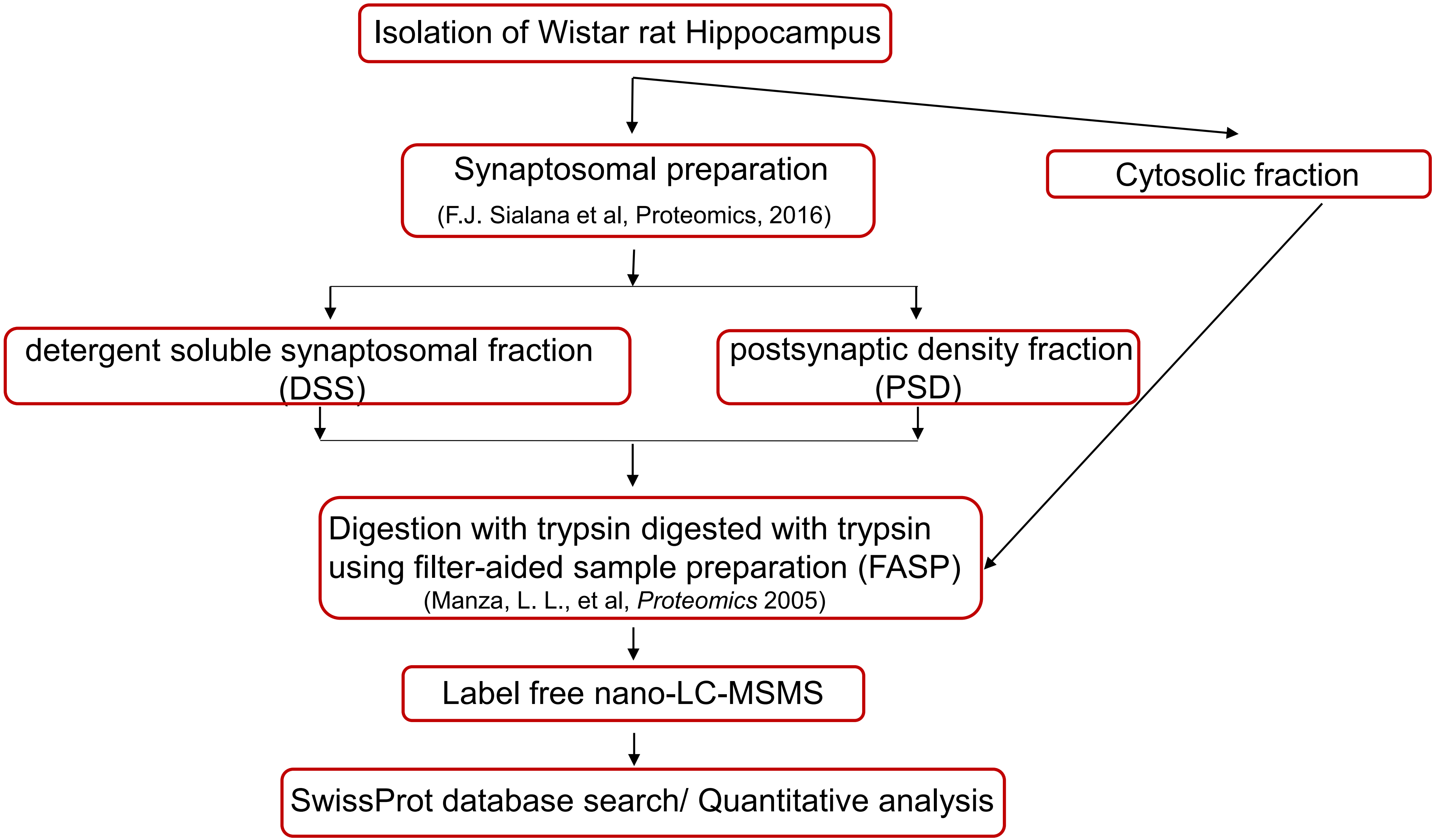

Isolation of Wistar rat Hippocampus
Synaptosomal preparation
(F.J. Sialana et al, Proteomics, 2016)
Cytosolic fraction
detergent soluble synaptosomal fraction (DSS)
postsynaptic density fraction (PSD)
Digestion with trypsin digested with trypsin using filter-aided sample preparation (FASP)
(Manza, L. L., et al, *Proteomics* 2005)
Label free nano-LC-MSMS
SwissProt database search/ Quantitative analysis

**Fig.7**

**a**

| Anti-Bbs5 antibody (mouse) | Anti-Bbs4 antibody (mouse) |
|---|---|
| Smith TS et al., Cell Mol Life Sci, 2013 | Prieto-Echagüe et al., Sci Rep, 2017 |
| Liew GM, et al., Dev.Cell, 2014 | Struchtrup A et al., Autophagy, 2018 |
| Sinha S et al., Inv Ophthalmol Vis Sci, 2014 | |
| Eguether T et al., Dev. Cell, 2014 | |
| Siller SS et al., Cell Cycle, 2015 | |
| Ye F et al., J Cell Biol, 2018 | |

**b**

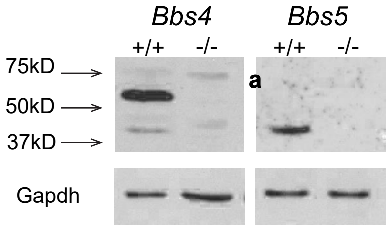

#### Supplementary Table 1

| Brain Region | Publication | Protein | Accession | Description | Coverage | # Peptides | # PSMs |
|---|---|---|---|---|---|---|---|
| Dorsal Striatum, rat | Sialana et.al., 2018 | BBS1 | D4A4U2 | Bardet-Biedl syndrome 1 homolog (Human) | 4 | 2 | 10 |
| synaptosomes | | BBS2 | Q99MH9 | Bardet-Biedl syndrome 2 protein homolog | 7 | 4 | 7 |
| TMT-labelling | | BBS4 | D4A8B1 | Bardet-Biedl syndrome 4 homolog (Human) | 2 | 1 | 2 |
| peptide prefractionation | | BBS5 | B2RZ48 | Bardet-Biedl syndrome 5 (Human) | 3 | 1 | 1 |
| | | BBS7 | Q66H90 | Bardet-Biedl syndrome 7 | 11 | 6 | 12 |
| | | BBS9 | F1M285 | Protein Bbs9 | 6 | 3 | 4 |
| Cortex, rat | Sialana et.al., 2016 | BBS10 | B1H296 | Bardet-Biedl syndrome 10 (Human) | 5 | 3 | 3 |
| synaptosomes | | BBS7 | Q66H90 | Bardet-Biedl syndrome 7 | 4 | 3 | 3 |
| TMT-labelling | | BBS2 | Q99MH9 | Bardet-Biedl syndrome 2 | 14 | 11 | 14 |
| peptide prefractionation | | | | | | | |
| Dentate gyrus, rat | Smidak et.al., 2017 | BBS1 | D4A4U2 | Bardet-Biedl syndrome 1 homolog (Human) | 11 | 4 | 6 |
| crude synaptosomes | | BBS2 | Q99MH9 | Bardet-Biedl syndrome 2 protein homolog | 10 | 7 | 12 |
| TMT-labelling | | BBS7 | Q66H90 | Bardet-Biedl syndrome 7 | 8 | 4 | 5 |
| peptide prefractionation | | BBS9 | F1M285 | Protein Bbs9 | 8 | 4 | 7 |
| Hippocampus, rat | this study | BBS1 | D4A4U2 | Bardet-Biedl syndrome 1 | 5 | 2 | 9 |
| synaptosomal fractions | | BBS2 | Q99MH9 | Bardet-Biedl syndrome 2 | 2 | 1 | 4 |
| label free analyses | | BBS4 | D4A8B1 | Bardet-Biedl syndrome 4 | 5 | 1 | 1 |
| single-shot 4hr gradient | | BBS5 | B2RZ48 | Bardet-Biedl syndrome 5 | 3 | 1 | 6 |
| | | BBS7 | Q66H90 | Bardet-Biedl syndrome 7 protein homolog | 5 | 3 | 5 |
| | | BBS9 | F1M285 | Bardet-Biedl syndrome 9 | 4 | 2 | 8 |
